# Complex benthic habitats retain larvae sinking in response to soluble cues: field study of coral reefs in wave-driven flow

**DOI:** 10.64898/2026.03.25.714321

**Authors:** M. A. R. Koehl, Michael G. Hadfield

## Abstract

Many benthic marine invertebrates disperse by releasing microscopic larvae carried by ocean currents to new sites, where they must settle into appropriate habitats and metamorphose to recruit. Species whose larvae settle in response to water-borne chemical cues live in topographically complex habitats. To study whether sinking in response to dissolved cues affects retention of larvae within complex habitats exposed to ambient water flow moving faster than larvae sink, we used the reef-dwelling sea slug, *Phestilla sibogae*, whose competent larvae stop swimming and sink in response to dissolved cue from their prey coral, *Porites compressa*. We conducted field experiments where dye-labelled water, neutrally buoyant particles, and larval mimics (particles that sank at the velocity of larvae of *P. sibogae*) were released together upstream of reefs of branching corals to determine if larval sinking in water above and within a reef affects larval retention within the reef. Wave-driven water flow measured above a reef in the field had instantaneous velocities peaking at 0.3 m s^-1^, driving slow net advection of water shoreward at ∼0.02 m s^-1^. Much slower wave-driven flow moved through the interstices within the reef. In this field flow, sinking by larval mimics caused their retention within a reef after dye-labelled water and neutrally buoyant particles had left. Such retention of sinking larvae within topographically complex benthic communities enhances successful recruitment by exposing larvae to high concentrations of cue for long periods, allowing them time to sink to surfaces, adhere, and undergo metamorphosis.

## 1. INTRODUCTION

Many benthic marine invertebrates disperse to new habitats by releasing microscopic larvae that are carried by ocean currents. After spending requisite times in the plankton developing the capacity to settle to the benthos and metamorphose into juveniles (defined as developmental competence, Hadfield et al. 2001), larvae must be transported near to surfaces by ambient flow to successfully recruit into suitable habitats (reviewed by Scheltema 1986, Koehl 2007, 2024) and must be able to choose where to settle from a variety of habitats across which they may be carried. Because submerged marine habitats are exposed to turbulent water flow, larval recognition of the “best spots” for attachment, metamorphosis, nutrition, growth, and reproduction with conspecific individuals must typically be a rapid process (i.e., the larva must settle in a suitable spot before it is carried away over less salubrious sites; Hadfield 2000).

### 1.1. Settlement behavior of larvae of benthic invertebrates in response to chemical cues

With data rapidly accumulating on the factors that induce competent larvae to settle into specific sites, a generality has emerged that most benthic invertebrate larvae attach to substrata in response to surface-bound chemical cues, most of which come from surface bacterial films (see Hadfield et al. 2025 for a broad review of this topic). However, there are also many examples of larval types from different phyla (e.g., cnidarians, bryozoans, annelids, molluscs, crustaceans, and ascidians) that are stimulated to settle in response to water-born chemical cues (reviewed by Hadfield & Paul 2001).

Invertebrate species whose larvae are sensitive to soluble cues appear to live in habitats with sufficiently 3-dimensional topographic complexity to slow water flow within them (e.g., seagrass beds, macroalgal assemblages, oyster beds, coral reefs). If water moving through the spaces within complex marine communities is slowed, then the concentrations of dissolved chemical cues from the benthos can increase, and this cue-laden water can remain within the community for significant periods of time (e.g., Hadfield & Koehl 2004, Koehl & Hadfield 2004). Furthermore, larvae carried into the slowly moving water within a complex habitat may not only encounter high concentrations of dissolved cue, but may also have longer residence times in the habitat to detect the cue and respond to it by descending and attaching to the substratum. Ambient water currents are slowed as they move through beds of seagrass or macroalgae (reviewed by Koehl 2022, 2023). In these complex macrophyte habitats, the larvae of such disparate organisms as the Queen Conch*, Aliger* (*Strombus*) *gigas* (Boettcher and Targett, 1998), the blue-crab *Callinectes sapidus* (Welch et al., 1997), and the sea urchin *Heliocidaris purpurescens* (Swanson et al. 2004) respond to dissolved chemical cues. Similarly, water is slowed as it moves through the interstices in oyster beds (Whitman & Reidenbach 2012), and larvae of the oyster *Crassostrea virginica* settle in response to a dissolved cue (Turner et al. 1994). Another important category of marine communities with complex topographies that slow the flow of water within them are coral reefs (e.g., Koehl & Hadfield 2004, Lowe et al. 2005a, 2005b, Monismith 2007, 2014, Reidenbach et al. 2009, Ascher & Shavit 2019, Webber & Huppert 2020, 2021, Davis et al. 2021, Pomeroy et al. 2023). The coralivorous sea slug *Phestilla sibogae* is an example of a reef-dwelling animal whose competent veliger larvae respond to a water-borne chemical cue from their prey coral, *Porites compressa*, by stopping swimming and sinking (Hadfield 1977, 1978, Hadfield & Pennington 1990, Hadfield & Koehl 2004), adhering to surfaces (Hadfield, 1978, Koehl & Hadfield 2004), and undergoing metamorphosis into benthic slugs (Hadfield & Scheuer 1985, Hadfield & Pennington 1990).

*P. sibogae* is the focus of the research described here. We have previously determined that water collected in the field from the spaces between coral branches within reefs dominated by *P. compressa* contains concentrations of the dissolved inductive substance in sufficiently high concentrations to induce the larvae of *P. sibogae* to respond as described above (Hadfield & Scheuer 1985, Hadfield & Koehl 2004, Koehl & Hadfield 2004). The goal of the present study is to determine in the field whether the sinking behavior of the larvae of *P. sibogae* can affect their retention in the cue-laden water within coral reefs.

### 1.2. Water flow across and through coral reefs can affect larval settlement

Ambient water flow near benthic habitats affects the settlement of larvae into those sites by a number of mechanisms, including delivering larvae, dispersing chemical cues, and washing settled larvae off surfaces (e g., reviewed by Koehl, 2007). *P. sibogae* is abundant on coral reefs in Kāneʻohe Bay, HI, where the large-scale (kilometers) water transport through the Bay has been determined (Bathen 1968, Lowe et al. 2009), and the smaller-scale flow velocities encountered by coral reefs have been studied. These reefs are composed mainly of branching corals, with *P. compressa* being one of the dominant species (e.g., Bahr et al. 2015).

The water velocities experienced by coral reefs in Kāneʻohe Bay have been measured in the field on the scale of cm’s to m’s (Koehl & Hadfield 2004, 2010, Lowe et al. 2005c), and this flow has been replicated and studied over skeletons of *P. compressa* in a wave flume on the scale of 0.1 mm’s to m’s (Koehl & Reidenbach 2007, Koehl et al. 2007, Reidenbach at al. 2007, 2008). We summarize the results of those studies here. The reefs in Kāneʻohe Bay are exposed to turbulent wave-driven water flow. Water in waves moves in vertical orbits, so the reefs experience back-and-forth shoreward-seaward flow with high instantaneous velocities (peaks of order 15 to 40 cm s^-1^, Koehl & Hadfield 2004, 2010, Lowe et al. 2005c, Koehl and Reidenbach 2007) producing slow net transport of water shoreward above the reefs (moving at rates of only 1 to 4 cm s^-1^, Koehl & Hadfield 2004, 2010, Lowe et al. 2009), and up-and-down water motion with slow net transport (rates of ∼1.5 cm s^-1^) up out of regions of a reef with convex or flat surfaces, and down into the reef in areas with concave depressions (Koehl & Hadfield 2010). In these porous reefs, composed mainly the branching coral *P. compressa,* water flowing through the interstices in the reef is also wave-driven, but both instantaneous shoreward-seaward water velocities (peaks of 2 to 4 cm s^-1^) and net shoreward water transport rates (∼0.5 cm s^-1^) are much slower than above the reef (Koehl & Hadfield 2004, 2010). These measurements for reefs in Kāneʻohe Bay are consistent with the results of other hydrodynamic models, flume studies, and field measurements of reef flow, which have shown that there is slow water movement through reefs, but that reefs exposed to waves have less flow reduction and greater vertical transport of water than do reefs exposed to unidirectional currents (Lowe et al. 2005a, Lowe et al. 2005b, Lowe et al. 2005c, Lowe et al. 2008, Monismith 2007, Lowe & Falter 2015, Stocking et al. 2016, Duvall et al. 2019, Davis et al. 2021, Pomeroy et al. 2023). When rough reef surfaces are exposed to wave-driven flow, a steep velocity gradient develops in the water within a few cm above the reef surface (Koehl & Reidenbach 2007), which spreads out the cue-laden water that is transported up out of the reef (unpublished field observations of dispersal of dye injected into water within reefs, Koehl & Hadfield). Other hydrodynamic studies of coral reefs have also noted such dispersion of water-borne materials just above the surface of a reef (Lowe et al. 2008, Stocking et al. 2016, Duvall et al. 2019, Ascher & Shavit 2019, Davis et al. 2021, Pomeroy et al. 2023). Planar laser induced fluorescence (PLIF) measurements in wave flume studies revealed that on the spatial scale of microscopic larvae, the chemical cue from the reef in the water above the reef surface is distributed in fine filaments swirling in odor-free water (Hadfield & Koehl 2004, Koehl & Reidenbach2007, Koehl et al. 2007, Reidenbach at al. 2007, 2008). Larvae swimming in that wave-driven turbulent flow above a reef have on-off encounters with cue as they swim or sink through the filaments, and their frequency of cue encounters increases if they move closer to the reef surface (Hadfield & Koehl 2004, Koehl et al. 2007). When exposed briefly to dissolved cue above a threshold concentration in a filament, larvae of *P. sibogae* stop swimming and then resume swimming when they exit the filament, with lag times of ∼1 s between entering or exiting a filament and the behavioral response (Hadfield & Koehl 2004). Encounters by larvae with filaments of cue above threshold concentration are so frequent in the water 10-20 cm above a reef (Koehl at al. 2007) that the larvae there should sink almost continuously.

An agent-based model of the transport of larvae of *P. sibogae* that stop swimming and sink when in filaments of dissolved coral cue in turbulent, wavy flow predicted that more larvae should settle into the upstream seaward-facing region of a reef than into downstream areas of the reef (Koehl et al. 2007). A study of the actual recruitment of *P. sibogae* onto reefs in Kāneʻohe Bay showed exactly this spatial pattern for newly settled *P. sibogae* (Hadfield et al. 2006).

Not only does ambient water flow affect the dispersal of chemical cues from and the delivery of larvae to benthic habitats, but it also can wash settled larvae off surfaces. To assess whether larvae that have landed on a reef are likely to be swept away by ambient water flow, the adhesive strengths of newly settled larvae of *P. sibogae* were measured (Koehl & Hadfield 2004). In a wave-flume study, a laser Doppler anemometer (LDA) was used to measure water velocities 200 μm (the height of larvae of *P. sibogae* on surfaces) from the tips of coral branches at the top of a *P. compressa* reef and from the surfaces of coral branches down within the reef (Reidenbach et al. 2007). Use of those velocities to calculate the hydrodynamic forces on the larvae revealed that forces on larvae on branch tips are greater than their adhesive strength, whereas those on larvae within the reef are lower than their adhesive strength. This result indicates that only those larvae that settle onto surfaces within a reef are likely to recruit successfully.

### 1.3. Objectives of this study

We conducted field experiments using larval mimic particles, which sink at the same velocity as do competent larvae of *P. sibogae* when exposed to a waterborne chemical cue from *P. compressa*, to determine if sinking in response to chemical cues in the water above and within a reef composed of branching corals affects larval retention in the water inside the reef structure.

The specific questions we addressed were:

1. How does water move above and through reefs composed of branching corals?
2. Do sinking larvae travel differently from the water that flows above or through such coral reefs?
3. Do sinking larvae travel differently from non-sinking, neutrally buoyant particles when carried in the water flowing above or within a coral reef?
4. Are there spatial patterns of the locations (at the top or within the fore reef, mid reef, and back reef) where most of the sinking larvae are retained, and do those patterns correlate with spatial patterns of larval recruitment into those reef locations?

## 2. MATERIALS AND METHODS

To better understand events involving settlement of larvae in the field, we conducted field experiments to determine whether induced sinking by larvae affects their transport above and through coral reefs by wave-driven turbulent ambient water flow. We measured the transport above and through coral reefs of water (marked by dye), of particles that sank at the same rate as competent larvae of *Phestilla sibogae* when responding to dissolved settlement cue from *Porites compressa* (“larval mimics”), and of neutrally buoyant particles.

### 2.1. Field sites

Water velocities and transport were measured over and through coral reefs in Kāneʻohe Bay on the island of Oahu, HI (N 21j27V, W 157j47V). These were the same reef sites, dominated by the coral *P. compressa*, described in Koehl and Hadfield (2004), Hadfield et al. (2007), and Koehl and Reidenbach (2007). Large ocean swells break on the reef crest across the open eastern side of the Bay, so smaller waves drive water movement across the patch reefs within the Bay (Bathen 1968, Lowe et al. 2005, 2009). Our sites were in the middle of the Bay where net flow across the reef was always in the shoreward direction, regardless of time in the tidal cycle (Bathen 1968, Lowe et al., 2009).

Transport experiments were conducted at three different reef sites on a given day. A total of 12 experiments were run. Mean wind speeds on the days of the experiments ranged between 4.5 and 6.7 ms^-1^, with gusts of 8.9 to 11.2 ms^-1^ (Kāneʻohe Bay MCAS Airfield Station https://www.wunderground.com/history/daily/us/hi/kaneohe/PHNG). Experiments were conducted during incoming or outgoing tides when the mean water depth 5 m seaward (upstream) of the edge of a reef was 0.87 m (SD = 0.2, n = 12 experiments). Wave heights, which were determined 5 m seaward of the fore reef immediately before an experiment by measuring the highest and lowest positions (to the nearest 0.01 m) of the water surface on a vertical pole for five successive waves, ranged between 0.05 and 0.20 m.

### 2.2. Measurement of water velocities above and within coral reefs

During one of the days when transport experiments were run, we also measured water velocities above and within a reef at a position midway between the upstream fore-reef and downstream back-reef edges (“mid reef” in Fig. 1, A). Water velocities 0.27 m above the reef were measured with a Marsh-McBirney Model 511 Electromagnetic Water Velocity Meter, which has a spherical probe (diameter = 3.81 cm; diameter of area sampled 11.4 cm). Simultaneous measurements of water velocities in the spaces within the reef (0.25 m below the top surface of the reef) were made with a Marsh-McBirney Model 523 Electromagnetic Water Velocity Meter, which has a smaller spherical probe (probe diameter = 1.27 cm, diameter of area sampled 3.8 cm). Both probes were oriented to measure the horizontal component of velocity in the seaward-shoreward direction, and the vertical component of velocity. Each probe was held in position by a stiff aluminum rod (diameter = 1.3 cm) supported by a scaffolding positioned so that it did not interfere with the water flow measured by the probe.

**Figure 1.**
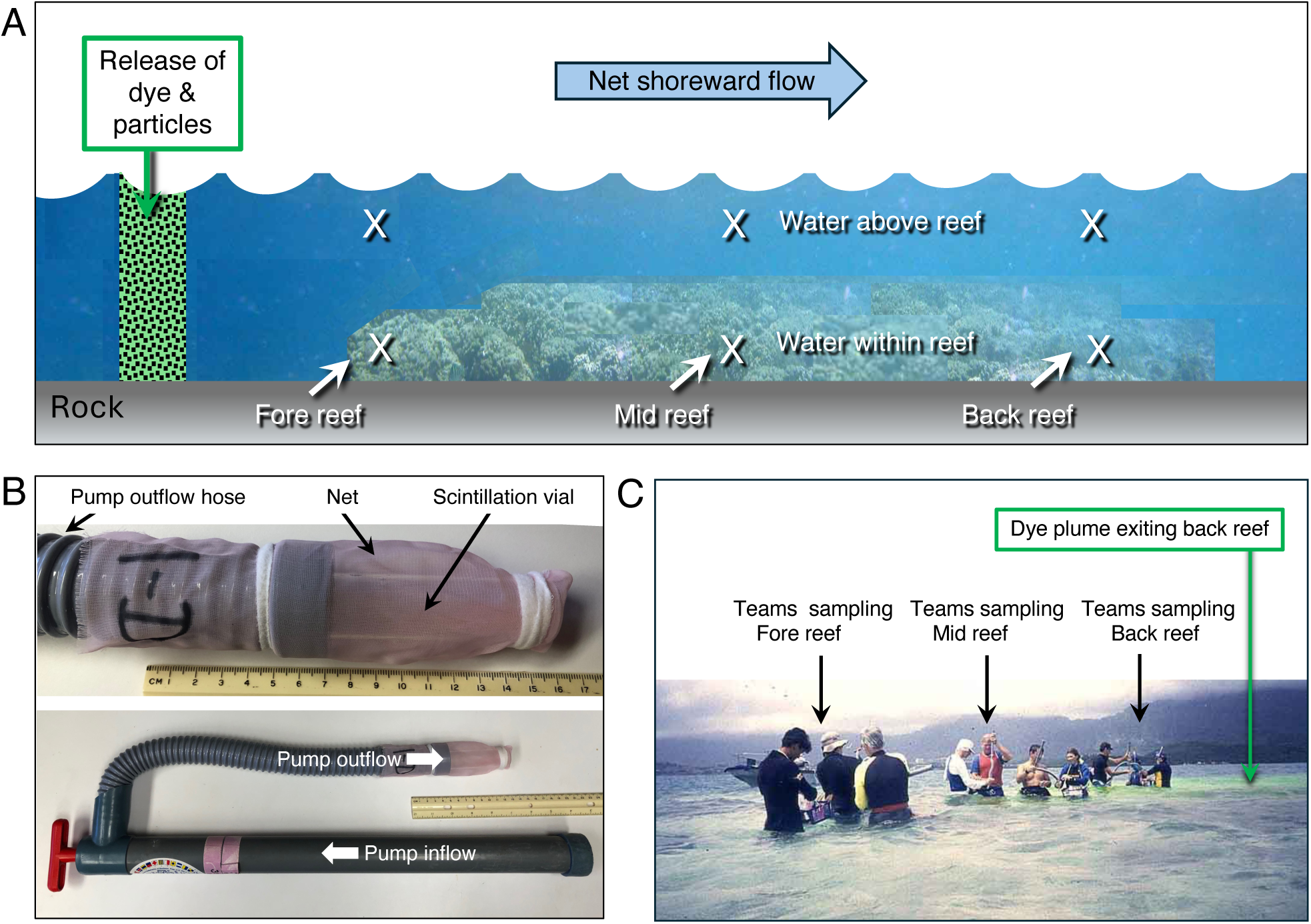
Field transport experiments. (**A)** Diagram of the positions of the release of fluoresceine-labelled water, larval mimics, and neutrally buoyant particles relative to a reef, and of the positions above and within a reef where water samples were taken (X’s). Blue arrow shows the direction of net shoreward flow of the wave-driven water motion across the reef. The wavy interface between the blue water and white air indicates the surface of the water. (**B)** Water samples were taken by hand-operated bilge pumps (lower image). Sampled water was passed through a net attached to the end of the outflow hose, and a water sample was caught in a scintillation vial inside the net (upper image). Pumps were held vertically when sampling. Numbers on ruler indicate cm’s. (**C)** Sampling teams during a field trial, viewed from a position seaward of the fore reef position, and to the right of the release point. The dye plume can be seen around the mid and back reef teams and exiting the back of the reef.

Cables from the probes were run to the Marsh-McBirney meters on a boat anchored nearby. The velocity data were converted from analog to digital using a DAQ-Card 1200 controlled by Labview 5 software (National Instruments, Austin, TX), and recorded by a Texas Instruments (Dallas, TX) TM 5000 laptop computer. The data acquisition rate was 60 Hz, and the time constant for the Marsh-McBirney meters was 0.2 s. The power spectral density of each velocity record was calculated by Welch’s method (Welch 1967, Rabiner & Gold 1975) using Matlab Signal Processing Toolbox 4.3 software.

### 2.3. Materials released for field transport experiments

For each field transport experiment (a field trial), a suspension of larval mimics (62.4 g) and neutrally buoyant particles (30.0 g) was made in 250 ml of a solution of fluoresceine (120.0 g l^-1^) in filtered (0.22 μm) sea water (“FSW”). Particles and fluoresceine were weighed to the nearest 0.1 g using a Mettler AL 100 Balance.

#### 2.3.1. Larval mimics

“Larval mimic” particles (Ultrafine Prisma Glitter #EA 6487, Capri Arts and Crafts) had sinking velocities similar to those of the larvae of *P. sibogae* when sinking in response to settlement cue from *P. compressa* (Hadfield & Koehl 2004), and that could easily be distinguished from natural particulate matter. The sinking velocities of 15 pieces of each color of Prisma Glitter that we used were measured using the procedure described in Butman (1988). There was no significant difference between the sinking velocities of different colors (ANOVA, p > 0.05). The mean of the mean sinking velocities of each color was 0.27 cm s^-1^ (SD = 0.02, n = 5 colors). The sinking velocities of larval mimics fell within the range of downward velocities (0.05 to 0.34 cm s^-1^) of competent larvae of *P. sibogae* that stop swimming when exposed to dissolved settlement cue from *P. compressa* (Hadfield & Koehl 2004).

Glitter particles had irregular shapes, so we measured the narrowest the width of individual larval mimic particles to the nearest 2.5 μm using an eyepiece micrometer in a Zeiss compound microscope (20x objective, 10x eyepieces). Measurements were done for 5 different colors of glitter. Glitter of a given color was dusted onto a piece of clear tape, and the first 5 isolated glitter particles observed were measured. There was no significant difference in width between the different colors (ANOVA, p > 0.05). The mean of the mean widths of larval mimics was 259 μm (SD = 11, n = 5 colors), which is close to the shell length of the larvae of *P. sibogae* (240 μm, Bonar & Hadfield 1974).

#### 2.3.2. Neutrally buoyant particles

The cysts of *Artemia salina* cysts (Brine Shrimp Eggs, San Francisco Bay Brand) were employed as neutrally buoyant particles. A 50 ml volume of dried cysts was stirred into FSW in 1000 ml beakers and left unagitated for 24 hours. The floating cysts were then skimmed from the top of the water and discarded, the neutrally buoyant cysts suspended in the FSW were decanted into another 1000 ml beaker and retained, and the negatively buoyant cysts at the bottom of the beaker were discarded. FSW was stirred into the beaker containing the decanted FSW and cysts, refilling it to 1000 ml, and the beaker was left for another 24 hours, after which any floating cysts were removed, the FSW with the neutrally buoyant cysts was decanted, and any cysts at the bottom of the beaker were discarded. This process was repeated 3 times (72 hours), after which the decanted water containing the neutrally buoyant cysts was passed through a coffee filter (Melitta 12-cup basket coffee filter) to collect the cysts, which were air-dried on the filters, and then brushed off the filters into glass jars for storage. At least 3 hours before a field experiment, these dried cysts were weighed and added to the fluoresceine solution to be released in the field. The solutions were examined right before release to determine that the cysts were still neutrally buoyant. Because viable *A. salina* cysts hatch as nauplii in 18 to 48 hours (Dees 1961, Vanhaecke & Sorgeloos 1982, Lavens & Sorgeloos 1987, Sellami et al. 2020), the cysts released in the field after 72 hours in FSW were not viable, and any nauplii from hatched cysts that might have been present in our decanted FSW were killed by drying on the coffee filters. The neutrally buoyant cysts ranged in size from about 220 to 250 μm.

### 2.4. Field release and sampling of dye and particles

The release and sampling positions relative to a coral reef for the transport experiments are diagrammed in Fig. 1A. For each field trial, a 250 ml suspension of larval mimics and neutrally buoyant particles in FSW labelled with fluoresceine was released at a mean distance of 5.6 m (SD = 0.2, n = 12 experiments) from the seaward (upstream) edge of a reef. The mean of the water depths at the release positions (measured to the nearest 0.01 m on a vertical pole) was 0.87 m (SD = 0.20, n = 12 experiments). The suspension was shaken in a wide-mouth jar and then rapidly poured out just below the water surface while the water was stirred up and down by a diving fin to mix the suspension evenly across the depth of the water. The dye enabled us to see how evenly the released suspension was mixed vertically in the water column and to track the movement of the water across the reef past our sampling positions.

### 2.5. Water sampling system

Water samples were taken using hand bilge pumps (Thirsty Mate Hand Pump, model 124PF8, Beckson Marine Inc.), which had an intake pipe and an output hose (Fig. 1B). We used a graduated cylinder to make three replicate measurements of the volume of the sample of water taken by one stroke of each pump and found that volumes were repeatable to the nearest 0.01 liter. The mean volume sampled by the pumps during one stroke was 0.67 liters (SD = 0.02, n = 6 pumps). Three strokes of a pump were used to collect each water sample (2.01 liters) taken in the field transport experiments.

Water leaving the pump outflow hose was passed through a cylindrical net attached by an elastic band to the end of the hose (Fig. 1B) to capture the larval mimics and neutrally buoyant particles. Nets (17 x 7 cm) were made of Georgette #751 polyester fabric (thread spacing 230 μm). To ensure that particles could not escape through the seams, they were sealed with hot-melt glue. A screw-cap scintillation vial (30 ml) held within the net by an elastic band (Fig. 1B) captured a water sample from the outflowing water.

### 2.6. Water sampling procedure

Water samples were taken at three positions (fore reef, mid reef, back reef) in the water above a reef (10 cm below the water surface) and at those same three positions within the reef infrastructure (Fig. 1, A). The wave-driven flow produced net water motion in the shoreward direction. We determined the direction of net flow by observing the path of a pulse of fluoresceine dye released from a syringe (with no needle) at the fore reef position for a field trial. Then the mid reef and back reef positions were selected to be directly downstream from the fore reef position. The water height above the reef (distance between the highest point on reef surface and the water surface) and the distance below the top of the reef of the within-reef positions were measured to the nearest 0.01 m on a vertical rod, and the distances from the seaward fore reef position to the mid reef and back reef positions were measured to the nearest 0.1 m using a transect line (Table 1).

**Table 1.** Sampling positions.

| Position<br>(n = 12 runs) | Distance from<br>seaward edge<br>of reef<br>(m)<br>mean ± SD | Water height<br>above reef<br>(m)<br>mean ± SD | Distance below<br>reef top of<br>within-reef sample<br>(m)<br>mean ± SD |
| --- | --- | --- | --- |
| Fore reef | 0 | 0.55 ± 0.20 | 0.10 ± 0.04 |
| Mid reef | 5.4 ± 3.2 | 0.59 ± 0.20 | 0.18 ± 0.14 |
| Back reef | 10.7 ± 3.2 | 0.70 ± 0.32 | 0.14 ± 0.08 |

A team of two people sampled the water at each position (Fig. 1C). The two teams sampling the fore reef stood on the rock next to the reef. The teams sampling the mid and back reef positions stood in depressions in the reef where they did not step on living coral. One person (A) in each team operated the pump, and the other person (B) managed the net and water samples, which were organized in floating basket. The first sample (t = 0 minutes) was taken when the dye reached the fore reef sampling positions. Subsequent samples were taken at t = 2, 4, 6, 8, and 10 minutes. Sampling times were managed by a timekeeper who called out when each sample was to be taken simultaneously by all the teams.

To take a water sample, the pump was held vertically by person A with the mouth of the intake pipe at the sampling position in the water above or within the reef. Water was drawn into the pump isokinetically by making sure the travel of particles observed moving next to the mouth of the pipe and into the mouth were the same. After a sample was taken, person B held the vial upright in the air so no water was spilled from the vial, while person B raised the intake pipe up into the air and pumped the plunger six more times until all the water was evacuated from the pump. Then person B removed the net and vial from the pump, put a cap on the vial, and sealed the net and vial in a zip-lock bag. After the net was removed from the output hose, person A rinsed the pump using clean FSW from the Kewalo Marine Laboratoy’s seawater system. A new net-and-vial was then attached to the pump outflow by person B for the next sampling time. At the end of each run, all samples were put into an opaque black plastic bag for transport to the lab so that the fluoresceine would not fade. Samples in black bags were then stored in a cold room at 4°C until processed.

### 2.7. Processing and measuring samples

Each sample was processed within one day of collection. The water in the scintillation vial was poured through a coffee filter to remove the particles, and then the concentration of fluoresceine in the water sample was measured using a Turner Fluorometer (Model 10-AU-005-CE). The particles captured on the net and any that fell into the ziplock bag were rinsed into the same coffee filter using FSW from a wash bottle. The coffee filter was then spread flat and air-dried. A plastic Petri dish (diameter = 150 mm) with a 1.0 cm x 1.0 cm grid marked on it was laid on top of the filter, and all the larval mimics and neutrally-buoyant-particles on the filter were counted.

### 2.8. Statistical analyses

Data used in parametric statistical tests met the assumption of normality (Shapiro-Wilke test) and homogeneity of variance (Levene’s test). Shapiro-Wilke tests, Levene’s tests, and linear regressions were done using Statistics Kingdom Online Calculators (Statistics Kingdom 2017, https://www.statskingdom.com). One-way ANOVA with post hoc Tukey HSD tests were done using the Astasa Online Statistical Calculator (N. Vasavada, 2016, https://astatsa.com). Paired-T tests were done using Statology Paired Samples T-test Calculator (Z. Bobbitt 2020,https://www.statology.org/paired-samples-t-test-calculator/). Concentrations of fluoresceine, larval mimics, and neutrally buoyant particles that had been normalized to the peak concentration at a position in a field trial were log-transformed before calculating linear regressions of concentration decrease as a function of time.

## 3. RESULTS

### 3.1. Water flow above and within a coral reef

Our reef sites were exposed to wave-driven turbulent water flow. Water in waves moves in vertical orbits, so our recordings of the instantaneous horizontal components of water velocity above a reef show back-and-forth flow in the shoreward-seaward direction (red plots in Fig. 2A), and our simultaneous recordings of the instantaneous vertical velocities show up-down flow (red plots Fig. 2C). Although the instantaneous water velocities in the shoreward-seaward direction above the reef peaked at values between 0.2 to 0.3 m s^-1^, there was a net slow advection of water shoreward above our reef sites at mean velocities of about 0.02 m s^-1^. There was also much slower wave-driven water flow in the interstices within the reef (blue plots in Figs. 2AC).

**Fig. 2.**
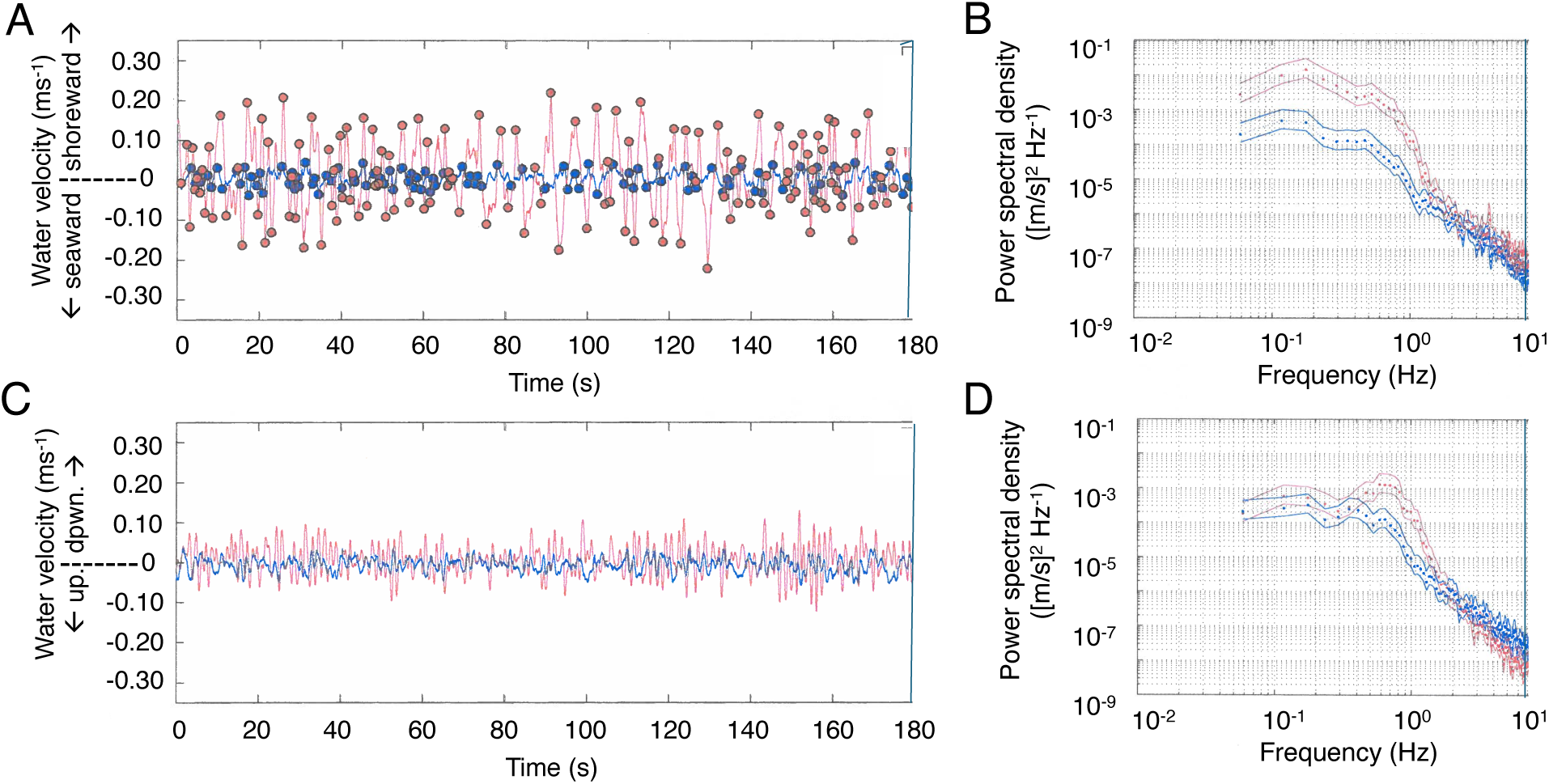
Water velocities recorded between field trials of transport experiments at the site of one of the field trials. Flow was recorded in the water 27 cm above a coral reef (red) and in the water in the interstices within a coral reef, 27 cm below the top of the reef (blue). (A) Shoreward-seaward components of velocity. (B) Turbulence spectra for shoreward-seaward flow. (C) Vertical components of velocities up out of the reef and down into the reef. (D) Turbulence spectra for vertical flow. In B and D, solid lines above and below the mean points indicate one standard deviation.

Turbulent eddies stir water and the materials it carries (e.g., larvae, dissolved chemical cues). By doing a spectral analysis of our velocity data, we plotted how much of the variation in velocity is due to fluctuations at different frequencies (Fig. 2BD). At frequencies below 1 Hz (large eddies and the waves, which had periods of 3 to 4 s), the power spectral density (PSD) of the water motion within the reef (blue in Fig. 2) was much lower than of the water flowing above the reef (red in Fig. 2). Moreover, the shoreward-seaward velocity fluctuations of the water above the reef showed high PSD both for waves and for larger eddies at lower frequencies (Fig 2B), whereas the vertical velocity fluctuations over the reef showed a PSD peak at the wave frequencies, but not for larger eddies (Fig. 2D). In contrast, at higher frequencies (i.e., smaller eddies), the PSD of the seaward-shoreward turbulence within the reef was similar to that above the reef (Fig. 2B), while the vertical turbulence within the reef was greater than above the reef at the scale of the smallest eddies we measured (Fig. 2D). These PSD data show that large-scale flow features were broken up when water entered the reef, but that the turbulent mixing of water by small eddies within the reef was as strong as above the reef.

### 3.2. Transport of water and water-borne particles above and through a coral reef

By sampling the water above and within the fore reef, mid reef, and back reef at timed intervals, we could track the transport of fluorescein-labelled water and the larval mimics and neutrally buoyant particles it carried above and within a reef. So that we could compare changes in concentrations of dye, larval mimics, and neutrally buoyant particles within a field trial, and so that we could compare field trials done on different days and reef sites, we normalized the concentrations of each sample. To do this, we divided the measured concentration of dye, of mimics, or of particles in each sample by the highest concentration (Concentration_max_) that was measured anywhere above or within the reef during that field trial of dye, mimics, or particles, respectively. An example of one of our twelve field trials is shown in Fig. 3 to illustrate the patterns we observe. These data show that water, larval mimics, and neutrally buoyant particles were carried across the reef more quickly in the rapidly flowing water above the reef than in the slowly moving water within the reef. Sinking larval mimics were retained in the water within the reef after the dye-labelled water and neutrally buoyant particles had left the reef.

**Fig. 3.**
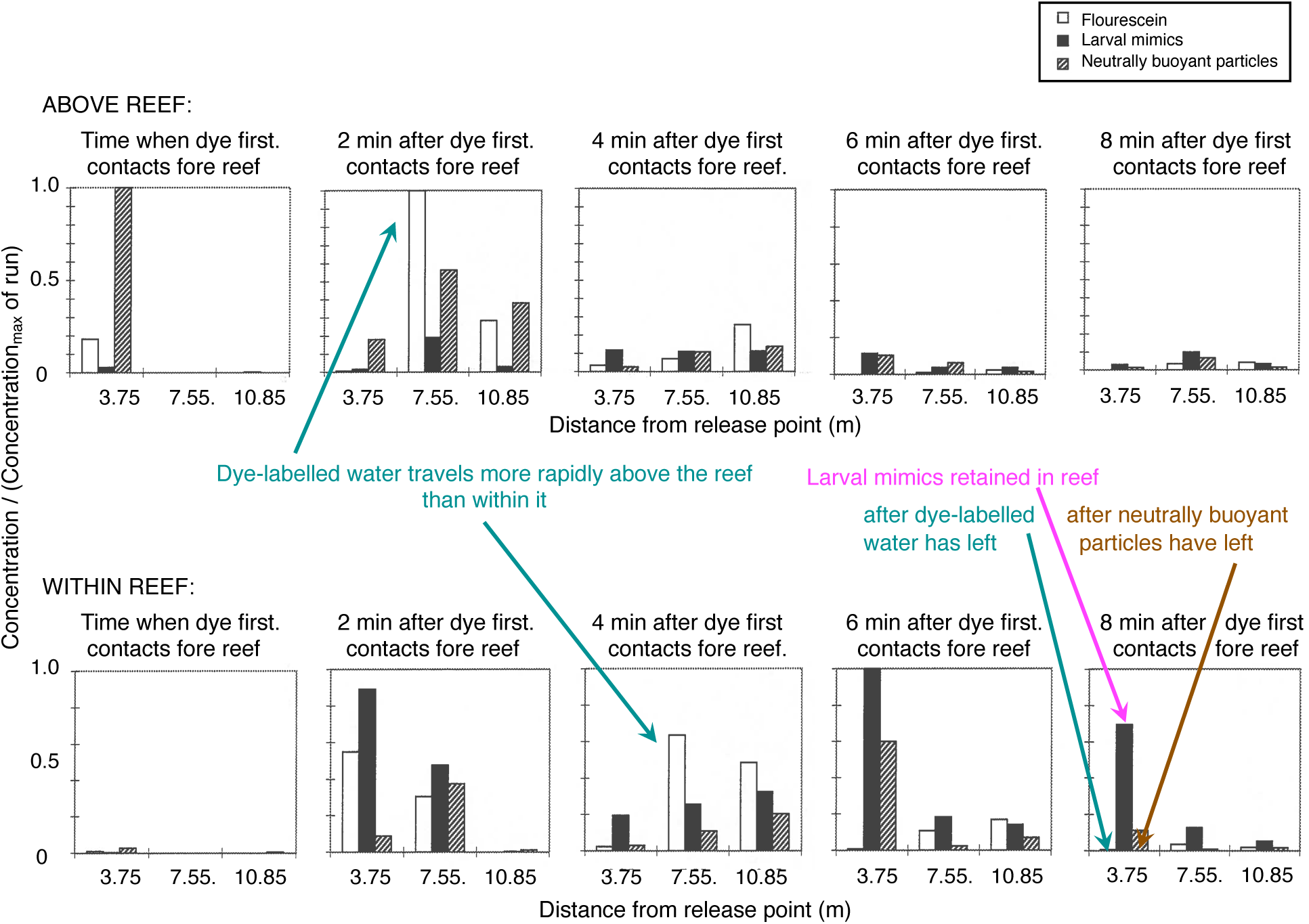
Example of concentrations of fluorescein, larval mimics, and neutrally buoyant particles measured in water samples taken during one field trial. Fluorescein concentrations (white columns) were normalized by dividing the measured concentration of a sample by the concentration of the sample taken during this field trial that had the highest fluorescein concentration (Concentration_max_) among all the samples taken at every location above and within the reef at every sampling time during the trial. Concentrations of larval mimics (black columns) and of neutrally buoyant particles (hatched columns) were normalized in the same way. Each graph shows the normalized concentrations measured at a given time, where t = 0 min is the time when the dye plume first contacts the fore reef. The x-axis of each graph indicates the distance of the sampling position from the release point seaward of the reef. The top row of graphs shows concentrations measured in the water flowing above the reef, and the bottom row of graphs shows concentrations measured in the water within the reef. Dye-labelled water travels more quickly above the reef than within it (for example, peak dye concentrations reach the mid reef position after 2 min above the reef, but after 4 minutes within the reef). At the end of the field trial, dyed water has left all the positions within the reef, but larval mimics remain.

### 3.3. Loss rates of larval mimics and neutrally buoyant particles from the water within a reef

To test whether sinking (as competent larvae of *Phestilla sibogae* do when exposed to dissolved chemical cue from the coral *Porites compressa*) is a mechanism that enhances larval retention in a reef, we determined whether the loss rates of sinking larval mimics from the water above or within a reef were lower than the loss rates of neutrally buoyant particles.

Loss rates for each run were determined for each position above or within a reef by plotting the concentration of fluorescein, larval mimics. and neutrally buoyant particles as a function of time. At each position, concentration increased as the dye-labelled water, mimics, and neutrally buoyant particles were carried into that position, and then decreased as the dye, mimics, and particles were washed away. So that we could compare values for dye-labelled water, mimics, and particles within a run at a given position, and so that we could compare data from different runs, we normalized the concentrations. For example, we normalized the concentration of larval mimics in each sample taken at a given position in a run by dividing it by the highest concentration (Concentration_Highest_) of mimics measured at that position during that run. (Note that Concentration_Highest_, which is the highest concentration at a specific position within or above a reef, is different from Concentration_max_, which is the highest concentration measured anywhere in a field trial.) Concentrations of dye-labelled water and of neutrally buoyant particles were normalized to Concentration_Highest_ in the same way. In plots of normalized concentration as a function of time during the period when dye, mimics or particles were being washed away, steeper negative slopes indicate greater loss rates. Because those concentrations did not drop off linearly with time, we log-transformed the normalized concentrations so that the slope of [ln (Concentration / Concentration_Highest_)] plotted as a function of time was linear. Some examples of such plots are given in Fig. 4. We used linear regression to calculate the slopes of the portions of each of those plots during which larval mimics were being washed away, and during which neutrally buoyant particles were being washed away. Those slopes were the loss rates for the larval mimics and for the neutrally buoyant particles, respectively.

**Fig. 4.**
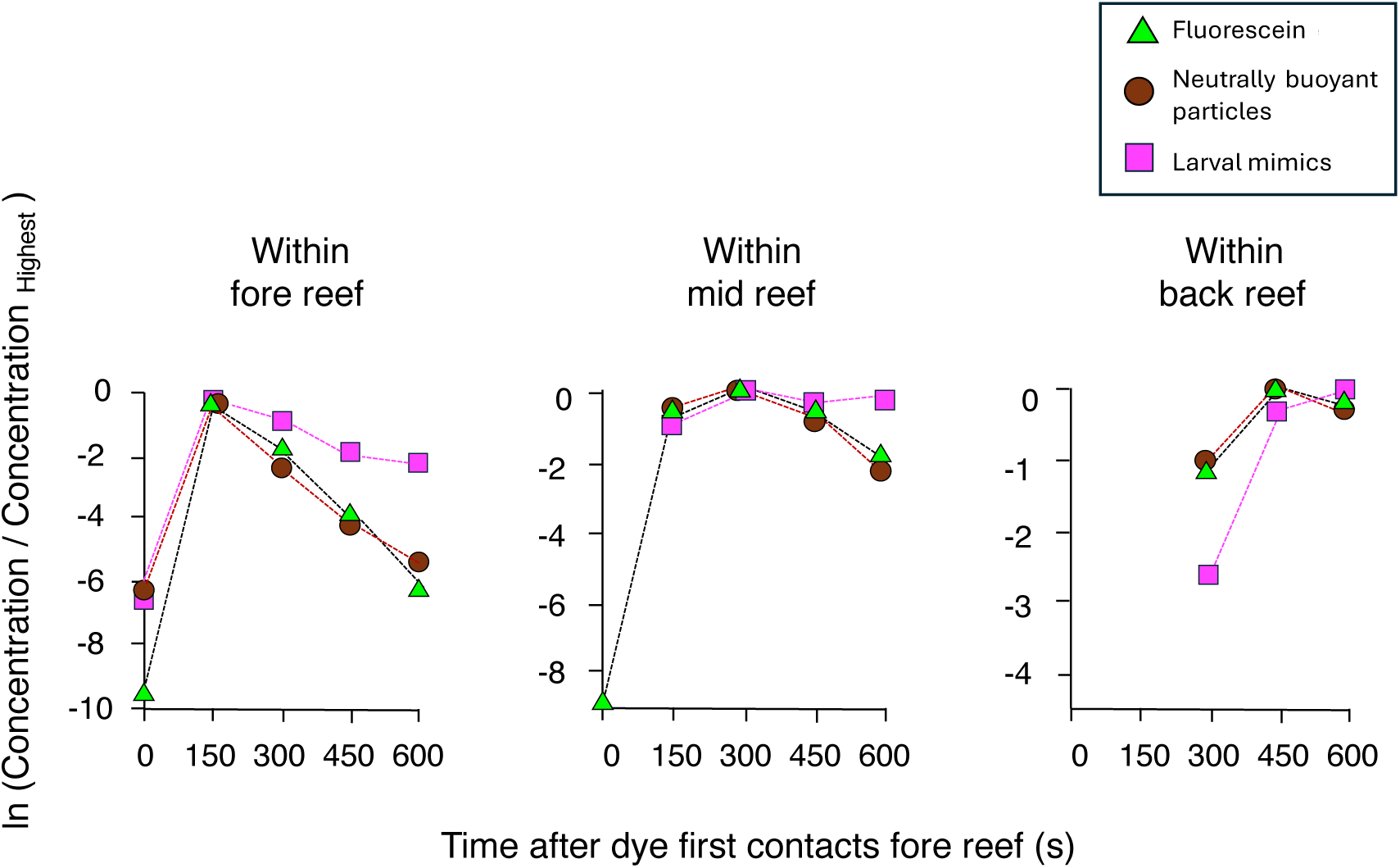
Concentrations of dye (green triangles), neutrally buoyant particles (brown circles) and larval mimics (pink squares) plotted as a function of time after a dye patch first touched the seaward edge of the fore reef. Within reef positions for one run are shown. Concentrations were normalized by dividing the concentration in a sample by the highest concentration achieved during that field trial at that position (Concentration_Highest_). These normalized concentrations were log-transformed so that linear regressions could be performed on the portions of the plots where concentration was declining. The slopes of those regressions are the loss rates reported in Table 2. Note that at the back reef position, the larval mimics were still accumulating at the end of this field trial while the fluorescein-labelled water and the neutrally buoyant particles were decreasing.

**Table 2.** Loss rates of larval mimics and neutrally buoyant particles. Data used in paired-t tests are the means of the loss rates for each position for all the field trials in which the loss rate regressions were calculated using ≥3 time points. We predicted that the loss rate of larval mimics would be lower than that of the neutrally buoyant particles, so we calculated the p for one-tailed paired t-tests (p < 0.05 for significance).

| Reef position | Above reef: |  | Within reef: |  |
| --- | --- | --- | --- | --- |
| | Mean loss rate ($s^{-1}$) of larval mimics | Mean loss rate ($s^{-1}$) Neutrally buoyant particles | Mean loss rate ($s^{-1}$) Larval mimics | Mean loss rate ($s^{-1}$) Neutrally buoyant particles |
| Fore | -0.45 | -0.44 | -0.39 | -0.54 |
| Mid | -0.22 | -0.43 | -0.33 | -0.63 |
| Back | -0.29 | -0.63 | -0.3 | -0.46 |
| mean = | -0.32 | -0.5 | **-0.34** | **-0.54** |
| SD = | 0.12 | 0.11 | 0.05 | 0.08 |
| n = | 3 positions | 3 positions | 3 positions | 3 positions |
| t (df) = | 1.76 (2) |  | 4.2 (2) |  |
| p = | 0.11 |  | 0.026 |  |

As illustrated by the examples in Fig. 4, the loss rates of fluorescein-labelled water and of neutrally buoyant particles from positions within the coral reef were similar, indicating that the neutrally buoyant particles and water travelled together. In contrast, the loss rates of larval mimics were slower (i.e., the slopes were shallower) in the fore reef and mid reef. In the example in Fig. 4, the concentration of mimics in the back reef had not yet started to decrease at the end of the experiment.

We tested whether the loss rates of sinking larval mimics were significantly lower than the loss rates of non-sinking neutrally buoyant particles in the water flowing above the reef and in the water flowing within the reef. For each reef position, we calculated the mean of the loss rates for all the field trials for which we had ≥3 time points during the period when concentrations were going down. Paired t-tests (Table 2) showed that there was no significant difference between the loss rates of larval mimics and neutrally buoyant particles in the rapidly moving turbulent water above the reef. However, in the slowly moving water within the reef, the loss rate of sinking larval mimics was significantly lower than the loss rate of non-sinking neutrally buoyant particles. This shows that sinking should enhance the retention of larvae within the interstices of a reef.

## 4. DISCUSSION

In the field research described here, we found that in turbulent wave-driven flow, water is transported shoreward slowly above coral reefs (at rates of a few cm s^-1^) and an order of magnitude more slowly within the interstices of those reefs. We also found that under these field conditions, passive sinking by larval mimics causes them to be retained in the slowly moving water within a coral reef, even though the mimics, like real larvae, sink more slowly than the instantaneous velocities of the wave-driven turbulent water flow that carries them through the environment.

### 4.1. Mass transport of water above and through reefs composed of branching corals

Our measurements of the transport of dye-labelled water above and through coral reefs in Kāneʻohe Bay corroborate predictions made on the basis of water velocity measurements at fixed points above or within reefs there (Koehl and Hadfield 2004, 2010). Although seaward-shoreward water velocities measured at points above reefs in those studies and in the present study showed the back-and-forth flow due to waves (with peak velocities of 15 to 40 cm s^-1^; Fig. 2A), the mean of those velocities over several minutes revealed a net shoreward water flow of only 1 to 4 cm s^-1^. It took 6 to 10 min for the dye-labeled patches of water moving above reefs to travel across them, from first arriving at the fore reef to last leaving the back reef (e.g., Fig. 2). These reefs were ∼11 m from fore reef to back reef (Table 1), so we estimated water transport rates by dividing 11 m by the duration we measured for each dye patch to move across a reef, We found that net shoreward transport rates of our dye-labelled water patches above reefs were 2 to 3 cm s^-1^, within the range of those calculated from mean velocities at points in the water above reefs (Koehl & Hadfield, 2004, 2010). Net shoreward water transport through reefs predicted by averaging velocity measurements at points within reefs yielded much slower transport rates (∼0.5 cm s^-1^, Koehl & Hadfield, 2010) than those above the reef. Although our instantaneous shoreward-seaward velocity measurements within reefs (Fig. 2A) were similar to those in the earlier studies, and the dye runs showed that water moved shoreward more slowly within the reefs than above them (e.g., Fig. 3), many of our dye transport field trial did not last long enough for all of the dye-labelled water within a reef to exit the reef (e.g., Fig. 4), so we have not estimated net shoreward transport velocities from those data.

When wave-driven flow moves into a coral reef, the large eddies associated with the waves are broken up, and smaller eddies are generated by the wakes that form around the individual coral branches within the reef (Monismith 2007, Hata et al. 2017, Ascher and Shavit 2019, Duvall et al. 2019, Davis et al. 2021, Pomeroy et al. 2023). We observed this pattern in our reefs. For turbulent, temporally fluctuating water flow, plots of power spectral density as a function of frequency (Fig. 2B,D) show how much power there is in eddies of different sizes (the higher the frequency, the smaller the eddy). In Figs. 2B (horizontal flow) and 2D (vertical flow), the red above-reef spectra show a peak at the wave frequencies (as was also seen in spectra of flow measured above reefs in Panama, Stocking et al. 2016), while the blue within-reef spectra do not have such wave peaks. In contrast, the power spectral density of the smaller-scale turbulent eddies at higher frequencies was the same for the water moving within our reefs as for the water flowing above those reefs (Fig. 2B,D), This indicates that turbulent mixing by the smaller eddies within the reef is as strong as it is above the reef. In addition, flow between coral branches is slower if branches are closer together (Chang et al. 2009, Chamberlain & Graus 1975, Reidenbach et al. 2006, Monismith 2007), so if there is variability in the widths of gaps between structural elements within a real reef in the field, this structural complexity can lead to meandering water paths and high spatial variability in water flow through the reef (Chang et al. 2009, Asher & Shavit 2019, Duvall et al. 2019, Davis et al. 2021). The small turbulent eddies within reefs exposed to wave-driven water flow mix the water and the materials it carries within the reef, both horizontally and vertically (Stocking et al. 2016, Davis et al. 2021, Pomeroy et al. 2023). That mixing enhances exchange of dissolved substances (such as chemical cues) between coral surfaces and the water in the reef (Lowe et al. 2005b, 2005c, Monismith 2007, Lowe et al. 2008, Duvall et al. 2019, Davis et al. 2021, Pomeroy et al. 2023). The mixing of water within a reef also enhances the delivery of water-borne particles and zooplankton (such as larvae) to within-reef surfaces (Reidenbach et al. 2009, Lowe & Falter 2015, Hata et al. 2017, Duvall et al. 2019).

### 4.2. The larval life of *Phestilla sibogae*

The behavior of different larval stages of *P. sibogae* can affect how ambient water flow helps their escape from or retention within coral reefs. The larvae of *P. sibogae* recruit very specifically to the surfaces of reefs where their prey coral, *Porites compressa*, is abundant. and they spend the remainder of their lives there (Harris 1975, Hadfield 1976). Feeding on the coral, the juvenile slugs grow to maturity, mate with others of their species, and lay their gelatinous egg ribbons on dead coral surfaces deep within the coral heads (Hadfield, personal observations over many years). Prehatching development within the egg ribbons requires about 7 days at 25-26 °C (Miller & Hadfield 1986). At hatching, these larvae are not yet competent to metamorphose, and they are extremely photopositive. Typically, they swim directly toward the major light source in straight paths, which will be directly upwards toward the reef surface in the field. This activity will thus remove the precompetent veligers from the coral and place them in rapidly moving water above the reef, which can transport them away from the reef. The larvae do not become competent to metamorphose in response to the settlement/metamorphosis inducer for about three days (Miller & Hadfield 1986), by which time they will surely have been carried away from the immediate corals where they were produced. However, as noted by Hadfield et al. (2006), they are unlikely to be carried out of Kāneʻohe Bay, due to its circulation patterns (Bathen 1968). By three days post-thatching, the larvae also become photo-indifferent and thus are likely to be vertically scattered by the wave-driven turbulent mixing in the waters overlying the coral reefs.

Those light-insensitive, competent larvae lower in the water column will thus approach each new reef at depths where they are either carried into corals or, if close enough to the top of the reef, can detect the settlement cue from the corals and respond by ceasing swimming and descending into the coral heads (Hadfield & Koehl, 2004). These are the metamorphically competent sinking larvae simulated by our larval mimics in the field studies reported here.

### 4.3. Sinking by competent larvae enhances their retention within coral reefs, increasing the likelihood of successful settlement

The results presented here demonstrate that under complex, real field conditions, slow sinking by larvae carried in turbulent wave-driven flow across coral reefs can enhance their retention within reefs. These field measurements verify predictions based on flume studies and agent-based modelling (Koehl et al. 2007) that sinking by larvae of *P. sibogae* in response to encounters with filaments of a dissolved chemical cue from a reef of *P. compressa* increases settlement onto the reef compared with larvae that do not sink. Further corroboration that the sinking behavior is responsible for larval retention in reefs is our discovery that the loss rate of sinking larval mimics from reef water in the field is significantly lower than that of non-sinking neutrally buoyant particles and of dyed water (Table 2, Fig. 4).

There are several reasons why being retained in the water within a coral reef should enhance the likelihood of successful settlement and metamorphosis by the larvae of *P. sibogae.* The slowly moving water travelling through the interstices of a reef dominated by *P. compressa* builds up concentrations of dissolved settlement cue high enough to induce the larvae to continue to sink, to attach to surfaces, and to undergo metamorphosis (Hadfield & Scheuer,1985, Hadfield & Koehl 2004, Koehl & Hadfield 2004). Larvae of *P. sibogae* retained within a reef are exposed to such cue concentrations for much longer durations than when riding in the water moving above a reef and thus are more likely to have enough time to land on and attach themselves to surfaces. When a competent larva of *P. sibogae* is induced to sink by the cue from *P. compressa*, its foot protrudes from its shell (Hadfield & Koehl 2004) and can stick to surfaces using mucus (Koehl & Hadfield 2004) produced in the propodial mucous gland (Bonar, 1974). However, larvae of *P. sibogae* require an exposure to cue of 1.5–2.0 h to fully develop their maximum adhesive strength to a substratum (Koehl & Hadfield 2004). In the slow flow within reefs, the hydrodynamic forces on settled larvae of *P. sibogae* are lower than their adhesive strength, whereas the forces they experience on coral branch tips at the top of a reef can rip them off those surfaces (Reidenbach at el. 2009). Thus, larvae that have settled and attached to surfaces within a reef are not likely to be washed away by ambient water motion and thus continue to be exposed to concentrations of cue high enough to induce metamorphosis, which requires about 6 hours of exposure to cue (Hadfield 1977). Another possible advantage of settling within reefs might be protection from the fish that prey on *P. sibogae.* These fish forage only on the exposed outer surfaces of corals, although crustacean predators on *P. sibogae* can move into the spaces within reefs to feed (Gochfeld & Aeby 1997).

### 4.4. Spatial patterns of retention of larval mimics and of recruitment by *P. sibogae*

The spatial patterns of retention of sinking larval mimics that we measured in the water within coral reefs in the field match the spatial patterns of recruitment of *P. sibogae* into reefs exposed to similar water flow. In a study of the recruitment of *P. sibogae* in Kāneʻohe Bay (Hadfield et al. 2006), the reefs monitored were in locations where, like the reefs we studied here, net water flow was always in the shoreward direction, regardless of the time in the tidal cycle (Bathen 1968). On those reefs, greater numbers of juvenile *P. sibogae* per kg of coral were found in the seaward fore-reef regions than in the mid-reef and back-reef areas (Hadfield et al. 2006), which is the same spatial pattern we found for larval mimics retained in reef water in the field (e.g., Fig. 3).

The spatial patterns of retention of larval mimics that we measured in the field are also consistent with the prediction of an agent-based model of larval transport in the turbulent flow above a coral reef (Koehl et al. 2007). The model predicted that if larvae of *P. sibogae* sink in response to encounters with filaments of cue from a reef, more of them should land on the fore reef than on the mid reef or back reef, which is the pattern of larval concentrations within the reef that we measured in this study.

Field experiments measuring where larval mimics first touch reef surfaces (Hadfield et al. 2006) showed the same spatial patterns that we found in the present study for the concentrations of larval mimics in reef water. In a prior study, Hadfield et al. (2006) attached sticky tabs that captured all larval mimics that contacted them to surfaces in different locations across coral reefs in the same part of Kāneʻohe Bay as our transport studies were done. They did field releases of larval mimics upstream of reefs and found that more mimics were retained on surfaces down within the reef than on coral branch tips at the top of the reef, and that more mimics landed on surfaces within the fore reef than landed on surfaces within the mid reef or back reef.

### 4.5. Conclusions

If competent larvae sink in response to encounters with dissolved chemical settlement cues, they can be retained in the slowly moving water in the interstices within coral reefs. While in the water within a reef, larvae can be exposed to steady concentrations of dissolved cue for settlement and metamorphosis, and larvae that have attached to surfaces are unlikely to be washed away by the slow ambient water flow. Thus, sinking by competent larvae in response to settlement cue can not only increase their probability of landing on a reef (Koehl et al. 2007), but also can enhance their likelihood of successful settlement and recruitment in the reef. This generality may apply to larvae of many other benthic invertebrates that respond to dissolved chemical cues in the water and whose habitats are sufficiently complex to retain those settlement cues, such as seagrass beds, kelp beds and oyster reefs (e.g., Turner et al. 1994, Boettcher & Targett 1998, Welch et al. 1997, Swanson et al. 2004, Reidenbach et al. 2013). Our study illustrates the importance of understanding how habitats with complex topography slow water transport and how larval responses affect recruitment, to inform the design of substrata for reef restoration projects, an area of active research in the global tropics.

## 5. ACKNOWLEDGEMENTS

The research reported here was financially supported by NSF grants OCE9907120 to MARK and OCE9907545 to MGH and Office of Naval Research grants N00014-95-1015 and N00014-95-1-1096 to MGH. The authors are extremely grateful to the group who made these field studies possible, most of them undergraduate students, graduate students, postdoctoral fellows and collaborating visitors in the Hadfield laboratory: Dr. A. C. Anil, Dr. Dmitri Boudko, Jason Boyd, Eugenio J. Carpizo-Ituarte, Kimberly Del Carmen, Dr. Eric Holm, Ella Meleshkevitch, Jeff Nagel, Brian Nedved, Dr. Valerie Paul, Scott Schellhammer, Dr. Robert Thacker and Catherine Unabia. Richard Chock, facilities manager for the Kewalo Marine Lab, cheerfully provided boat access to our field site, and Tim Cooper provided technical assistance on data analysis. This paper is publication no. XXXXX from the School of Ocean and Earth Sciences and Technology at the University of Hawaiʻi at Mānoa. The present work is available as preprint at https://www.biorxiv.org/.

## REFERENCES

Asher S, Shavit U (2019) The effect of water depth and internal geometry on the turbulent flow inside a coral reef. J Geophys Res: Oceans 124:3508–3522. doi10.1029/2018JC014331

Bahr KD, Jokiel PL, Toonen RJ (2015) The unnatural history of Kāneʻohe Bay: coral reef resilience in the face of centuries of anthropogenic impacts. PeerJ 3e950; doi:10.7717/peerj.950

Bathen KH (1968) A descriptive study of the physical oceanography of Kaneohe Bay, Oahu, Hawaii. Hawaii Institute of Marine Biology, Honolulu, HI. Tech Rep No. 14, 353 pp

Boettcher AA, Targett NM (1998) Role of chemical inducers in larval metamorphosis of queen conch, *Strombus gigas* Linnaeus: relationship to other marine invertebrate systems. Biol Bull 194:132–142

Bonar DB 1974. Metamorphosis of the marine gastropod *Phestilla sibogae* Bergh (Nudibranchia: Aeolidacea): I. Light and electron microscope analysis of larval metamorphic stages. J Exp Mar Biol Ecol 16:227–255

Bonar DB, Hadfield MG (1974) Metamorphosis of the marine gastropod *Phestilla sibogae* Bergh (Nudibranchia: Aeolidacea) I. Light and electron microscope analysis of larval and metamorphic stages. J Exp Mar Biol Ecol 16:227–255

Butman CA, Grassle JP, Buskey EJ (1988) Horizontal swimming and gravitational sinking of “Capitella sp.” (Annelida: Polychaeta) larvae: implications for settlement. Ophelia 29:43–57.

Chamberlain JA, Graus RR (1975) Water flow and hydromechanical adaptations of branched reef corals. Bull Mar Sci 25:112–125

Chang, S, Elkins, C, Alley M, Eaton J, Monismith S (2009) Flow Inside a Coral Colony Measured Using Magnetic Resonance Velocimetry. Limnol Oceanog 54:1819–1827

Davis KA, Pawlak G, Monismith SG (2021) Turbulence and coral reefs. Ann Rev Mar Sci 13: 343–373

Duvall MS, Hench JL, Rosman JH (2019) Collapsing complexity: quantifying multiscale properties of reef topography. J Geophys Res: Oceans 124:5021–5038

Hadfield MG, Carpizo-Ituarte EJ, del Carmen K, Nedved BT (2001) Metamorphic competence, a major adaptive convergence in marine invertebrate larvae. Amer Zool 41:1123–1131

Hadfield MG, Koehl MAR (2004) Rapid behavioral responses of an invertebrate larva to dissolved settlement cue. Biol Bull 207: 28–43

Hadfield MG, Faucci A, Koehl MAR (2006) Measuring recruitment of minute larvae in a complex field environment: The corallivorous nudibranch *Phestilla sibogae* (Bergh). J Exp Mar Biol Ecol 338: 57–72

Hadfield MG, VJ Paul (2001) Natural chemical cues for settlement and metamorphosis of marine invertebrate larvae. In: McClintock JB, Baker W (eds) Marine Chemical Ecology. CRC Press, Boca Raton, FL, p 431–461

Hadfield MG (1976) Molluscs associated with living tropical corals. Micronesica 12: 133–148

Hadfield, MG, Scheuer D (1985) Evidence for a soluble metamorphic inducer in *Phestilla*: ecological, chemical and biological data. Bull Mar Sci 37:556–566

Hata T, Madin JS, Cumbo VR, Denny M, Figueiredo J, Harii S, Thomas CJ, Baird AH (2017) Coral larvae are poor swimmers and require fine-scale reef structure to settle. Sci Rep 7:2249 doi:10.1038/s41598-017-02402-y

Kaandorp JA, Koopman EA, Sloot PM, Bak RP, Vermeij MJ, Lampmann LE 2003. Simulation and analysis of flow patterns around the scleractinian coral *Madracis mirabilis* (Duchassaing and Michelotti). Phil Trans R Soc Lond B 358:1551–57

Koehl MAR (2007) Hydrodynamics of larval settlement into fouling communities. Biofoul 23:357–368

Koehl MAR (2022) Ecological biomechanics of marine macrophytes. J Exper Bot 73:1104–1121 doi10.1093/jxb/erab536

Koehl MAR (2023) Of corpses, ghosts, and mirages: Biomechanical consequences of morphology depend on the environment. J Exp Biol 226: jeb245442. doi:10.1242/jeb.245442

Koehl MAR (2024) A life outside. Ann Rev Mar Sci 16:17.1–17.23. doi/abs/10.1146/annurev-marine-032223-014227

Koehl MAR, Cooper T (2015) Swimming in an unsteady world. Integr Comp Biol 55:683–697 doi:10.1093/icb/icv092

Koehl MAR, Hadfield MG (2004) Soluble settlement cue in slowly moving water within coral reefs induces larval adhesion to surfaces. J Mar Sys 49:75–88

Koehl MAR, Hadfield MG (2010) Hydrodynamics of larval settlement from a larva’s point of view. Integr Comp Biol 50:539–551

Koehl MAR, Strother JA, Reidenbach MA, Koseff JR, Hadfield MG (2007) Individual-based model of larval transport to coral reefs in turbulent, wave-driven flow: Effects of behavioral responses to dissolved settlement cues. Mar Ecol Prog Ser 335:1–18

Koehl MAR. Reidenbach, MA (2007) Swimming by microscopic organisms in ambient water flow. Exp Fluids 43: 55–768

Lavens P, Sorgeloos P (1987) The cryptobiotic state of Artemia cysts, its diapause deactivation and hatching: A review. In: Sorgeloos P, Bengtson P, Decleir DA, Jaspers W, (Eds) Artemia Research and its Applications Vol. 3. Ecology, Culturing, Use in Aquaculture. Universal Press, Wetteren, Belgium, p 27–63

Levenstein MA, Gysbers DJ, Marhaver KL, Kattom S, Tichy L, Quinlan Z, Tholen HM, Kelly LW, Vermeij MJA, Johnson AJW, Juarez G (2022) Millimeter-scale topography facilitates coral larval settlement in wave-driven oscillatory flow. PLoS One 17(9), e0274088, doi:10.1371/ʻjournal.pone.0274088

Lowe RJ, Koseff JR, Monismith SG. (2005a) Oscillatory flow through submerged canopies. 1. Velocity structure. J Geophys Res: Oceans 110 C10, doi:10.1029./2004.JC002788

Lowe RJ, Falter JL (2015) Oceanic Forcing of Coral Reefs. Ann Rev Mar. Sci. 7:43–66 doi: 10.1146/annurev-marine-010814-015834

Lowe RJ, Koseff JR, Monismith SG, Falter JL (2005b) Oscillatory flow through submerged canopies: 2. Canopy mass transfer. J Geophy Res: Oceans 110 C10, doi10.1029/2004JC002789

Lowe RJ, Shavit U, Falter JL, Koseff JR, Monismith SG. (2008) Modeling flow in coral communities with and without waves: a synthesis of porous media and canopy flow approaches. Limnol. Oceanogr 53:2668–2680

Lowe RJ, Falter JL, Monismith SG, Atkinson MJ (2009) Wave-driven circulation in a coastal reef-lagoon system. J Phys Oceanogr 39:869–89.

Lowe RJ, Falter JL, Bandet MD, Pawlak G, Atkinson MJ, Monismith SG, Koseff JR. 2005c. Spectral wave dissipation over a barrier coral reef. J Geophys Res: Oceans 110 C4, doi10.1029/2004JC002711

Miller SE, and Hadfield MG (1986) Ontogeny of phototaxis and metamorphic competence in larvae of the nudibranch *Phestilla sibogae* Bergh (Gastropoda: Opisthobranchia). J Exp Mar Biol Ecol 97:95–112

Monismith SG (2007). Hydrodynamics of Coral Reefs. Annu Rev Fluid Mech 39:37–55 doi:10.1146/annurev.fluid.38.050304.092125

Monismith SG (2014) Flow through a rough, shallow reef. Coral Reefs 33:99–104

Pomeroy AWM, Ghisalberti M, Peterson M, Farooji VE (2023) A framework to quantify flow through coral reefs of varying coral cover and morphology. PLoS ONE 18(1): e0279623. doi10.1371/journal.pone.0279623

Rabiner LR, Gold B (1975) Theory and Application of Digital Signal Processing. Prentice-Hall, Englewood Cliffs, NJ, 762 pp

Reidenbach MA, Koseff JR, Monismith SG (2007) Laboratory experiments of fine scale mixing and mass transport within a coral canopy. Physics of Fluids 19:75–107

Reidenbach MA, Koseff JR, Koehl MAR (2009) Hydrodynamic forces on larvae affect their settlement on coral reefs in turbulent, wave-driven flow. Limnol Oceanogr 54:318–30

Reidenbach MA, Koseff JR, Koehl MAR (2009) Hydrodynamic forces on larvae affect their settlement on coral reefs in turbulent, wave-driven flow. Limnol Oceanogr 54:318–330

Reidenbach MA, Koseff JR, Monismith SG, Steinbuck JV, Genin A (2006) The effects of waves and morphology on mass transfer within branched reef corals. Limnol Oceanogr 51:1134–1141

Sellami I, Naceur HB, Kacem A (2020) Study of cysts biometry and hatching percentage of the brine shrimp Artemia salina (Linnaeus, 1758) from the Sebkha of Sidi El Hani (Tunisia) according to successive generations. Aquacult Stud 21: 41–46

Stocking JB, Rippe JP, Reidenbach MA (2016) Structure and dynamics of turbulent boundary layer flow over healthy and algae-covered corals. Coral Reefs 35:1047–59

Swanson RL, Williamson JE, De Nys R, Kumar N, Bucknall MP, Steinberg PD (2004) Induction of settlement of larvae of the sea urchin *Holopneustes purpurascens* by histamine from a host alga. Biol Bull 206:161–72

Swearer SE, Treml EA, Shima JS (2019) A review of biophysical models of marine larval dispersal. Oceanog Mar Biol, 325–356

Turner EJ, Zimmer-Faust RK, Palmer MA, Luckenbach M, Pentchef ND (1994) Settlement of oyster (*Crassostrea virginica*) larvae: Effects of water flow and a water-soluble chemical cue. Limnol. Oceanog 39:1579–1593

Vanhaecke P, Sorgeloos P (1982) International study on Artemia XVIII. The hatching rate of Artemia cysts - A comparative study. Aquacul Eng 1:263–273

Webber JJ, Huppert HE (2020) Stokes drift in coral reefs with depth-varying permeability. Phil Trans R Soc A: Mathematical, Physical and Engineering Sciences 378(2179) doi10.1098/rsta.2019.0531

Webber JJ, Huppert HE (2021) Stokes drift through corals. Env Fluid Mech 21:1119–1135

Welch PD (1967) The use of fast Fourier transform for the estimation of power spectra: a method based on time averaging over short, modified periodiograms. IEEE Trans. Audio Electroacoust. 15:70–73

Welch JM, Rittschof D, Bullock TM, Forward Jr RB (1997) Effects of chemical cues on settlement behavior of blue crab *Callinectes sapidus* postlarvae. Mar Ecol Prog Ser 154:143–53

Whitman ER, Reidenbach MA (2012) Benthic flow environments affect recruitment of *Crassostrea virginica* larvae to an intertidal oyster reef. Mar Ecol Prog Ser 463:177–191

